# All-Assay-Max2 pQSAR: Activity predictions as accurate as 4-concentration IC_50_s for 8,558 Novartis assays

**DOI:** 10.1101/620864

**Authors:** Eric J Martin, Valery R Polyakov, Xiang-Wei Zhu, Prasenjit Mukherjee, Li Tian, Xin Liu

**Affiliations:** Novartis Institute for Biomedical Research, 5300 Chiron Way, Emeryville, California, 94608-2916, United States; China Novartis Institutes for BioMedical Research Co., Ltd., 2F, Building 4, Novartis Campus, No. 4218 Jinke Road, Zhangjiang, Pudong, Shanghai 201203, China

**Author notes:** Xin Liu: School of Biological Sciences, The University of Edinburgh, Alexander Crum Brown Road, Edinburgh, EH9 3FF, UK, Prasnjit Mukheerjee: Gilead Sciences, 333 Lakeside Drive, Foster City, California 94404, United States.

**Keywords:** pQSAR, PLS, reduced profile, ChEMBL, transfer learning, chance correlation, multi-task model, applicability domain

## Abstract

Profile-QSAR (pQSAR) is a massively multi-task, 2-step machine learning method with unprecedented scope, accuracy and applicability domain. In step one, a “profile” of conventional single-assay random forest regression (RFR) models are trained on a very large number of biochemical and cellular pIC_50_ assays using Morgan 2 sub-structural fingerprints as compound descriptors. In step two, a panel of PLS models are built using the profile of pIC_50_ predictions from those RFR models as compound descriptors. Hence the name. Previously described for a panel of 728 biochemical and cellular kinase assays, we have now built an enormous pQSAR from 11,805 diverse Novartis IC_50_ and EC_50_ assays. This large number of assays, and hence of compound descriptors for PLS, dictated reducing the profile by only including RFR models whose predictions correlate with the assay being modeled. The RFR and pQSAR models were evaluated with our “realistically novel” held-out test set whose median average similarity to the nearest training set member across the 11,805 assays was only 0.34, thus testing a realistically large applicability domain. For the 11,805 single-assay RFR models, the median correlation of prediction with experiment was only *R*^2^ _ext_=0.05, virtually random, and only 8% of the models achieved our standard success threshold of *R*^2^ _ext_=0.30. For pQSAR, the median correlation was *R*^2^ _ext_=0.53, comparable to 4-concentration experimental IC_50_s, and 72% of the models met our *R*^2^ _ext_>0.30 standard, totaling 8558 successful models. The successful models included assays from all of the 51 annotated target sub-classes, as well as 4196 phenotypic assays, indicating that pQSAR can be applied to virtually any disease area. Every month, all models are updated to include new measurements, and predictions are made for 5.5 million Novartis compounds, totaling 50 billion predictions. Common uses have included virtual screening, selectivity design, toxicity and promiscuity prediction, mechanism-of-action prediction, and others.

## INTRODUCTION

Developing accurate, predictive quantitative structure-activity relationship (QSAR) models with wide applicability domains, for most biological assays, has long been a goal of computational scientists. As computer power has increased, building QSAR models has become increasingly automated and robust.^1^ However, while conventional, single-task QSARs can give accurate predictions for compounds very like the training set, such as random held-out test sets, they generally fail on the far more novel compounds project teams actually order for testing from virtual screens. Massively multi-task QSAR models improve the accuracy and applicability domain.^2^ Additionally, studies^3, 4^ have introduced *in vitro* biological assay information as descriptors in QSAR. Recent studies^5-7^ employing experimental *in vivo* or *in vitro* test results as descriptors filled missing biological values using either pure chemical information ^6, 7^ or a mixture of chemical and biological data ^5^.

Profile-QSAR (pQSAR) is an automated, massively multi-task, QSAR method that employs predicted biological activities as descriptors. Originally developed over 10 years ago for kinases^8^, the 2-step pQSAR method was based on the hypothesis that the activity of a compound in a new kinase assay could be modeled as a linear combination of the compound’s activities in all previously studied kinase assays. First, conventional single-assay (SA) random forest regression (RFR) models were trained on available pIC_50_ data for 728 biochemical or cellular kinase assays. The full profile (FP) of 728 RFR predicted pIC_50_s for each compound was then used as the compound descriptor vector for a second partial least squares (PLS) regression model of each assay. Hence the name “Profile-QSAR”. Through this 2-step procedure, every compound and every experimental pIC_50_ from all 728 assays informs each new pIC_50_ prediction. This expanded the applicability domain of these PLS models to cover the entire Novartis (NVS) archive. The transfer-learning among kinase assays also dramatically increased the accuracy over SA models. Evaluation with a “realistically novel” held-out test set showed the median correlation with experiment improved from the nearly random *R*^2^_ext_ = 0.12 for the original SA RFR models to *R*^2^_ext_ = 0.55 for pQSAR, comparable to experimental 4-concentration IC_50_s^9^. Likewise, the number of models exceeding *R*^2^_ext_=0.30, our rule-of-thumb for successful virtual screening, increased from 20% to 80%. An additional advantage of pQSAR is that it does not require combining different assays for a given target. The PLS models are trained only on the assay(s) the project team is currently using, but all other assays for that target still inform the model with optimal scaling (along with all other assays in the profile). Subsequently, pQSAR models were also built for several other large families: proteases^10^, GPCRs, CYP450s, and other ATP-binding protein targets^11^.

Project teams are increasingly working on targets outside these large protein families. Furthermore, many cellular assays are purely phenotypic--not specific to any target or family. Thanks to improvements in the implementation, we were able to address this with a massive “all-assay” (AA) pQSAR combining all NVS IC_50_ assays. The success of the AA pQSAR demonstrated the effectiveness of transfer learning between protein families as well as phenotypic assays. However, using the much larger AA FP of RFR predictions as descriptors to train PLS models on assays with few pIC_50_s could lead to overfitting, so a variable selection method was required. Overzealous variable selection from such a large descriptor pool also invites chance correlations ^12^ which was proven to be minimal by Y-scrambling.

This report describes building a massive multi-task AA pQSAR combining 11,805 diverse NVS dose-response assays crossing all protein families as well as phenotypic assays. To facilitate public comparisons to other methods, a second massive AA pQSAR was built for 4276 diverse, publically available ChEMBL^13^ assays.

## METHODS

### Data sets

Two, large, multi-family datasets were used: publically available, dose-response activity assays from ChEMBL version 24.1, downloaded from https://www.ebi.ac.uk/chembl, last accessed June 22, 2018, and a larger internal dataset of proprietary NVS assays downloaded November 2018. In both cases, compound/activity pairs were retrieved for all assays reporting IC_50_, AC_50_, EC_50_, K_*i*_, K_*d*_, A_*max*_, and other quantitative fields. All such values are referred to here as “IC_50_s”. Only the largest molecule was retained from multi-structure records. Compounds were neutralized, tautomers were standardized, counter-ions were stripped, and all structures were converted to canonical SMILES through RDkit (v2018.09.1.0). Duplicates were removed within each assay. “Qualified” values below or above the assay detection limits were offset by 1 log unit and treated as quantitative. Where concentration units were reported, they were converted to molar, and negative log transformed to give pIC_50_s. Occasionally no units were given in ChEMBL. The units were then guessed at from the range of values, converted to molar, and labeled as “uncertain units”. While these assignments might be incorrect, correlations will not be affected, so it will not affect the contribution to the PLS models. Multiple pIC_50_s for a given compound were averaged, discarding qualified values if quantitative values were available. Only assays with at least 50 pIC_50_s with a standard deviation of at least 0.5 were retained. In both data sets, some assays appeared to be almost entirely included within other assays. If the correlation between overlapping members of two or more assays exceeded 99%, the smaller assay(s) were excluded from the profile in the second step of PLS modeling. The final NVS data included 11805 assays, 1.8 million compounds, and 18.3 million pIC_50_s. The final ChEMBL data included 4276 assays, 0.5 million compounds, and 1.4 million pIC_50_s. The distributions of assay size for NVS and ChEMBl are presented in Figure S1. The specific assay identifiers for 4276 ChEMBL assays with corresponding number of compounds, assay type, and target family are listed in Supplemental Table S1.

### Realistic training/test set splits

The final production models for virtual screening are trained on all the data at every step. However, evaluating the quality of a model requires a held-out test set. The “realistic training/test set split” was previously shown to mirror the novelty of the compounds that project teams ordered from actual virtual screens.^9^ The previously described algorithm was ported from KNIME and ICM to python. The compounds are first clustered. For smaller assays, a hierarchical clustering is employed. For assays with more than 10,000 compounds, the Butina clustering algorithm ^14^ is used instead. The training set is collected starting with the largest cluster, and proceeding to successively smaller clusters until 75% of the compounds have been gathered. The remaining singletons and small clusters make up the “realistic” test set. The box plots in Figure S2 summarize histograms of the Tanimoto similarity (Tc) between the each test set member and the nearest training set member for realistic and random splitting of NVS and ChEMBL compounds.

### pQSAR workflow

Figure 1 depicts the 2-step pQSAR 2.0 algorithm, as previously described for the kinase family.^9^ First, conventional, SA RFR QSARs are trained for each assay on all available sparse historical activity data using scikit-learn ^15^ RandomForestRegressor (v0.20.2). No test-set data are withheld in this step (see below). Parameters were defaults except the number of trees was set to 200. Compound descriptors are RDkit (v2018.09.1.0) Morgan radius 2 substructure fingerprints with 1024 bits. These models are used to predict (or fit) a full pIC_50_ activity matrix of all compounds by all assays. In the second step of “FP pQSAR”, these FPs of RFR predicted pIC_50_s are used as compound descriptors to train a PLS model for each assay, individually, using scikit-learn ^15^ PLSRegression (v0.20.2). For “reduced-profile” pQSAR, in order to reduce the compound’s descriptor vectors for the PLS models in the second step, the training set experimental pIC_50_s for the assay currently being modeled are compared to predictions from the RFR models for each of the other assays. An assay is excluded from the reduced profile if the correlation does not reach a threshold, indicated by the greyed out columns in Figure 1. Two models are built for each assay: with thresholds of *R*^2^>0.2 and *R*^2^>0.05. The higher threshold generally gives better models, but for some assays that leaves few or no assays in the profile. It is not clear how many assays in the profile are ideal, so for simplicity, the model with the better *R*^2^_ext_ on the test set is retained. We call this “max2”.

**Figure 1.**
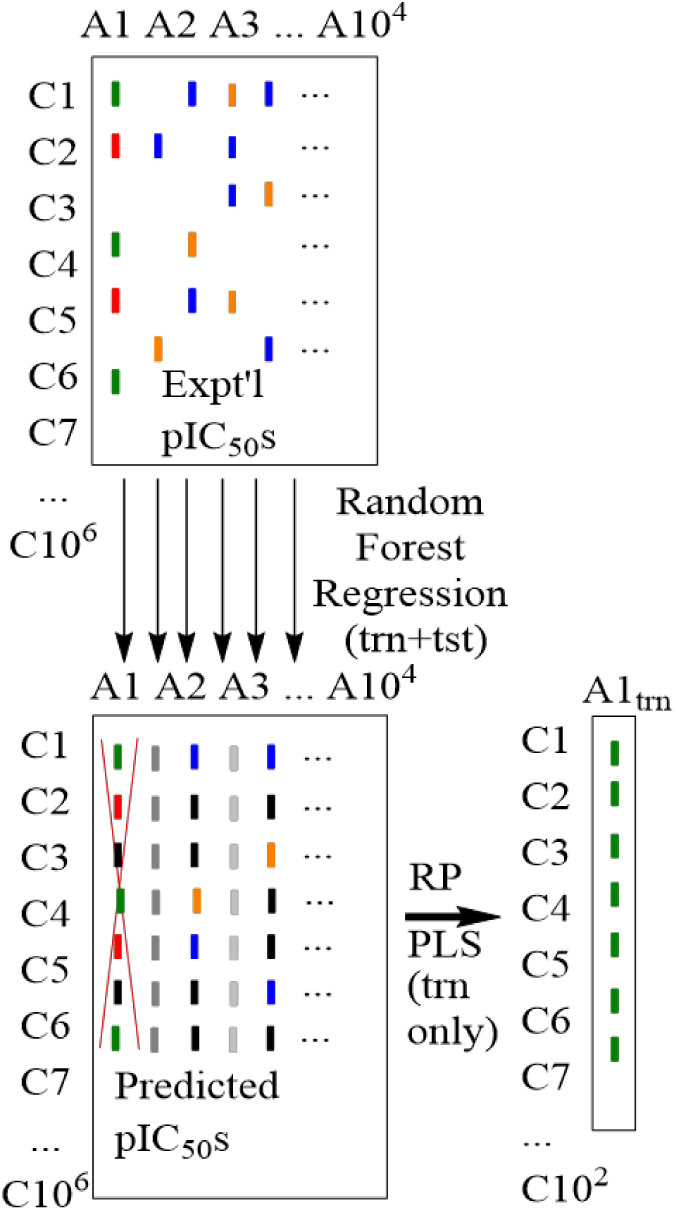
Reduced profile two-step pQSAR workflow focusing on training assay A1. (red = A1_test_, Green = A1_train_, Brown= A*_test_, Blue=A*_train_, Black = A*_NA_). The red X indicates that the assay being modeled is not included in the profile. Grey indicates columns excluded from the reduced profile because they do not correlate well with A1_train_.

When evaluating pQSAR, just the training set compounds from the realistic training/test set split are used to train the PLS model. The assay being modeled, as well as RFR models that correlate with R^2^>0.99 with the assay being modeled, are excluded from the profile as indicated by the red “X”. For predictions on new compounds, they are first run through the RFR models to generate the predicted pIC_50_ profile, which is then run through the final PLS model to predict the activity. Using models trained on each of the columns in the first RFR step, followed by a model based on the new row in the second PLS step, results in every experimental pIC_50_ from the reduced profile informing every prediction. A custom model for a new assay does not require training additional RFR models. It only requires making RFR predictions for any new training compounds to add new rows to the existing profile matrix, then training a new PLS model. Of course, this new assay will not contribute to improving the models for other assays in the panel. The entire process including retraining of all SA RFR models is performed monthly, and at that time the new assay will contribute to the other models as well.

### Y-scrambling

Y-scrambling ^16^ was used to estimate the number of apparently successful models that are actually due to chance correlations. Randomly reassigning the pIC_50_s among the structures for each assay were performed on both the NVS and ChEMBL data sets. SA RFR models were trained. As expected, few models had *R*^2^_ext_>0.3. Y-scrambled AA FP and AA max2 PLS Models were trained on these meaningless SA RFR “profiles”, and the number of “successful” models (*R*^2^_ext_>0.30) was counted and analyzed for comparison with models built on the unscrambled data. AA FP and AA max2 PLS models for the Y-scrambled assay data were also trained on the “real” SA RFR profiles.

## RESULTS AND DISCUSSION

A total of 11,805 dose-response assays measured on at least 50 unique compounds and with pIC_50_ standard deviation greater than 0.5 were found in the NVS database. The experimental data included 1.8 million unique compounds, and 18.3 million IC_50_ measurements. On average, each compound was measured on 10 assays and each assay was measured for 1550 compounds. The complete compound by assay matrix was only 0.1% complete. Timely pQSAR calculation on this large dataset required improvements in the implementation, including porting all KNIME, R and ICM code to python, although some infrastructure is still in C#. As seen from the bottom curve in Figure 2A, using the realistic training/test set split, 92% of these assays failed to produce successful (*R*^2^_ext_>0.30) SA RFR models, and the median correlation for SA RFR models was only *R*^2^_ext_=0.05, virtually random. The next curve up shows that applying the previously published pQSAR 2.0 algorithm using the FP dramatically improved the results, increasing the median correlation with experiment to *R*^2^_ext_=0.38 and 58% of the assays achieving successful models. The median across the 11,805 assays of the average Tanimoto similarity between each member of the test set and the nearest member of the training was only 0.34. Thus, this evaluation of model prediction quality covers a huge applicability domain, virtually the entire NVS archive. This is essential for estimating virtual screening success where chemical novelty is highly valued, and for poly-pharmacology prediction where the available training compounds for an off-target model would typically be unrelated to the current project. By contrast, the median average test set to training set similarity for the random held out tests sets was 0.62. Metrics based on this test set would be unreliable for estimating the accuracy of the models in most real drug discovery scenarios. The box plots in Figures S2A and S2B summarize, for the 11,805 NVS assays, histograms of test set to training set similarity for the realistic and random splits, respectively. We strongly recommend that the average test set similarity be reported whenever a performance metric is presented for a model, so the reader will understand the applicability domain for which the performance has been evaluated. This issue is well illustrated by the following ChEMBL performance summary.

**Figure 2.**
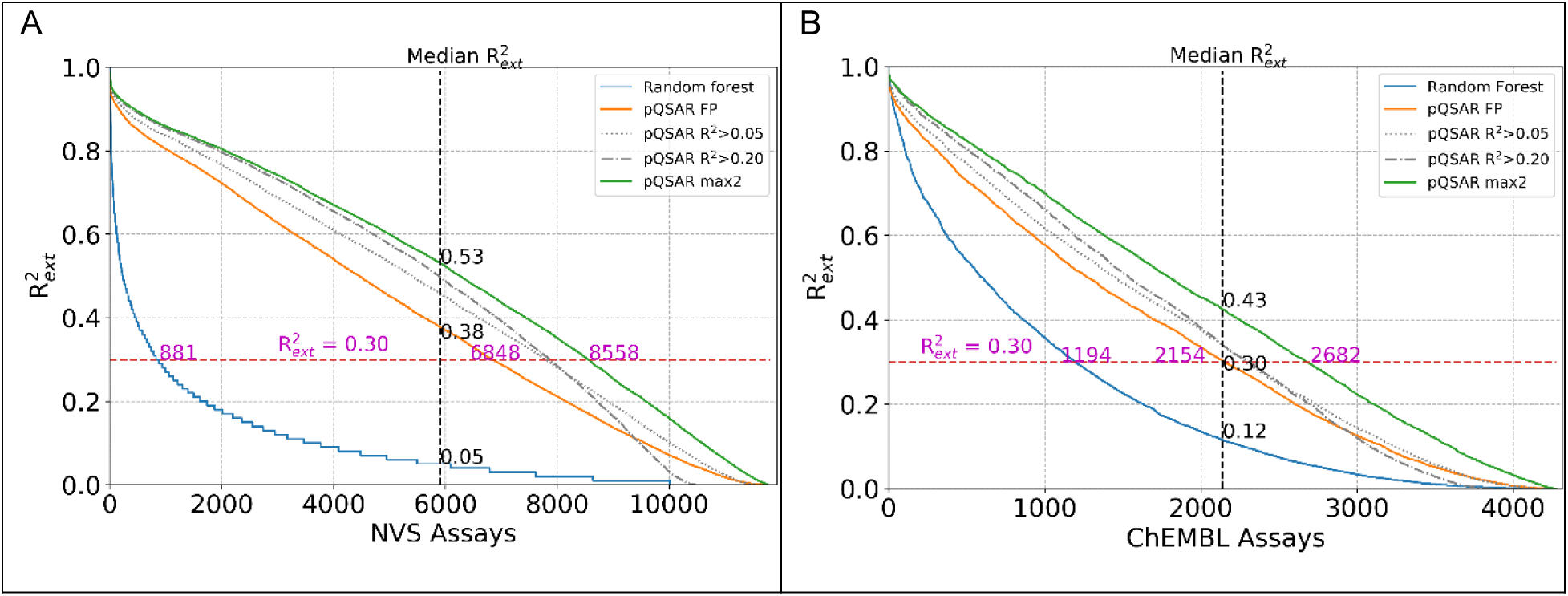
Correlations between prediction and experiment using the “realistic” training/test set split for SA RFR models and pQSAR models with full and reduced profiles: (A): NVS assays; (B): ChEMBL assays.

In order to present a non-proprietary analysis with full scientific reproducibility, the same criteria were employed to retrieve assays from the public ChEMBL database. The 4276 such assays, covering 500,000 compounds, included 1.4 million pIC_50_s, for a 0.1% complete matrix. However, the average compound was measured on only 2.8 assays and the average assay had only 327 compounds. Supplemental Table S2 lists ChEMBL (compound) ID, assay ID, observed pIC_50_s and max2 predicted pIC_50_s for the 1.4 million activity measurements. Supplemental Table S3 lists the RDkit canonical SMILES for the half million ChEMBL compounds. Figure 2B shows that 28% of these assays gave successful (*R*^2^_ext_>0.30) SA RFR models with a median correlation of *R*^2^_ext_=0.12, a much higher fraction than the NVS SA RFR models. This is partly due to more chance correlations (see below). However, this might mainly result from the much more homogeneous ChEMBL compound sets. The median across the 4276 assays of the average Tanimoto similarity between the realistic test and training sets was 0.63, much more similar than NVS. The median average random test set similarity is fully 0.77. This would seriously overestimate the performance for virtual screening where novelty is a compound selection criterion. The box plots in Figures S2C and S2D summarize, for the ChEMBL assays, histograms of test set to training set similarity for the realistic and random splits.

As Figure 2B shows, FP pQSAR also greatly improved the ChEMBL results, although not as dramatically as in the NVS case, increasing the median correlation with experiment to *R*^2^_ext_=0.30 and 50% of the assays achieving successful models.

It is known that PLS models lose accuracy with increasing numbers of irrelevant descriptors.^17^ With 11,805 NVS SA RFR models in the AA FP as compound descriptors, but an average of 1,162 and sometimes as few as 37 training IC_50_s, the ratio of independent variables to observations reduces the performance even of a powerful latent-variable regression method like PLS. Methods to identify unhelpful RFR models and build reduced profile pQSARs were therefore explored. For simplicity, methods that would employ the same subset for all PLS models were tried first. Predictions were not improved by using only SA RFR models trained on assays exceeding a minimal number of IC_50_s, or setting a minimum SA RFR *R*^2^_ext_.

The earlier family-specific (FS) pQSARs can be thought of as an intermediate case of reducing the profile, where the FP of SA RFR predictions from just a specific family was used for all pQSAR models from that family. Overall, these FS-FP pQSARs were equal or better, sometimes much better, than AA FP pQSARs. Yet, only one third of the assays are members of large families. A more general method to reduce the profile was therefore sought that would also apply to the remaining two thirds of the assays from phenotypic screens or smaller families.

Methods that employed a tailored, reduced profile for each assay were therefore explored. An attractive feature of pQSAR is simplicity. It is a turnkey method with virtually no architecture or hyper-parameter retraining. Furthermore, well-known variable selection methods would be frustrated by the unique “realistic” training/test set split. *I.e.* hyper-parameters such as variable selections optimized against random subsets of the training set might not be optimal for the far more novel realistic test set. Finally, we wanted a simple method that ideally would decrease the compute resources of the second PLS step, by using smaller profiles, rather than increasing it with a laborious variable selection algorithm. An effective compromise was to test in advance of PLS, and only retain for each PLS model, the RFR models in the profile whose predicted pIC_50_s correlated with the experimental pIC_50_s in the dependent assay’s training set. Higher thresholds generally gave better predictions, but some PLS models failed because there were few or no RFR models meeting the threshold. A simple compromise was to build PLS models using 2 thresholds: *R*^2^>0.2 and *R*^2^>0.05, and use whichever gave the better final *R*^2^_ext_. We call this “max2”. As Figure 2A shows, the AA max2 pQSAR median *R*^2^_ext_ was 0.53 on the realistic test set, now again comparable to high-throughput 4-concentration IC_50_s. 72% of the assays now met our success criterion. For the ChEMBL data, the AA max2 pQSAR median *R*^2^_ext_ was lower at 0.43, with 63% of the assays meeting the success criterion as shown in Figure 2B. The predicted ChEMBL AA max2 pQSAR pIC_50_s are listed in Supplementary Table S2.

For AA FP pQSAR, using the linear PLS regression method worked better than using the non-linear RFR in the second step, as had been previously found for kinases. To further test this result, recursive random forest regression (RRFR) ^18^, a method that combines variable reduction with RFR, was compared to max2 PLS. Figure S3 shows that the median *R*^2^_ext_ for NVS RRFR pQSAR is 0.51 with 62% of models having *R*^2^_ext_ > 0.3, worse than max2 PLS despite the far more expensive variable reduction and non-linear regression calculation. ChEMBL RRFR pQSAR also lags max2 PLS, with median *R*^2^_ext_ = 0.29 and 49% of models with *R*^2^_ext_ > 0.3. This is at variance with conclusions drawn by many other studies ^19, 20^. This unusual result might be due to the larger extrapolations required by the realistic test set compared to the more usual random test sets.

Table 1 compares the accuracy of the AA FP pQSAR models with the 3 methods for reducing profile size for the larger assay families: AA-max2, FS-FP, FS-max2. For ChEMBL, the order in almost every case is the same: AA-max2 ≥ FS-max2 ≫ FS-FP > AA-FP, both in the median *R*^2^_ext_ and in the number of models with *R*^2^_ext_ > 0.30. The only exception is ChEMBL ADME models (which may not even count as a family), where AA-FP is better than FS-FP. For NVS, AA-max2 and FS-max2 are very close. AA-max2 produces more successful models in most cases, but FS-max2 produces more for GPCRs and NHRs. We do not consider the differences large enough to justify maintaining separate family-specific models.

**Table 1.** Model performance of full profile (FP) and best of reduced profile (max2) on all assays or family-specific ones.

| | | | Median $R^2_{\text{ext}}$ | | | | % $R^2_{\text{ext}} > 0.3$ | | | |
| --- | --- | --- | --- | --- | --- | --- | --- | --- | --- | --- |
|  | Family | Assay count | AA-max2<sup>ab</sup> | FS-max2<sup>c</sup> | FS-FP<sup>d</sup> | AA-FP | AA-max2 | FS-max2 | FS-FP | AA-FP |
| ChEMBL | Binding | 2523 | 0.55 | 0.40 | 0.26 | 0.26 | 75% | 58% | 46% | 46% |
|  | Phenotypic | 2299 | 0.57 | 0.43 | 0.28 | 0.28 | 75% | 62% | 48% | 48% |
|  | Functional | 1689 | 0.59 | 0.49 | 0.35 | 0.32 | 78% | 64% | 54% | 52% |
|  | GPCR | 685 | 0.54 | 0.41 | 0.28 | 0.26 | 76% | 61% | 48% | 45% |
|  | Kinase | 665 | 0.56 | 0.50 | 0.42 | 0.35 | 79% | 69% | 59% | 56% |
|  | Proteinase | 289 | 0.58 | 0.48 | 0.33 | 0.30 | 78% | 67% | 53% | 50% |
|  | Ion channel | 215 | 0.60 | 0.41 | 0.30 | 0.24 | 77% | 63% | 51% | 47% |
|  | NHR | 123 | 0.44 | 0.26 | 0.21 | 0.14 | 67% | 46% | 41% | 35% |
|  | ADME | 55 | 0.46 | 0.13 | 0.08 | 0.20 | 69% | 31% | 24% | 42% |
| NVS | Functional | 8070 | 0.57 | 0.56 | 0.42 | 0.41 | 74% | 73% | 61% | 61% |
|  | Phenotypic | 5506 | 0.60 | 0.59 | 0.47 | 0.44 | 76% | 74% | 65% | 63% |
|  | Binding | 3446 | 0.47 | 0.47 | 0.37 | 0.32 | 70% | 69% | 59% | 52% |
|  | Kinase | 2407 | 0.62 | 0.63 | 0.55 | 0.50 | 85% | 85% | 78% | 73% |
|  | Enzyme | 1085 | 0.39 | 0.38 | 0.32 | 0.24 | 61% | 60% | 53% | 42% |
|  | GPCR | 815 | 0.34 | 0.37 | 0.29 | 0.20 | 55% | 57% | 49% | 36% |
|  | Protease | 698 | 0.46 | 0.48 | 0.43 | 0.32 | 68% | 68% | 61% | 52% |
|  | Toxicity | 289 | 0.44 | 0.39 | 0.36 | 0.30 | 69% | 60% | 55% | 51% |
|  | Ion channel | 249 | 0.23 | 0.20 | 0.16 | 0.13 | 41% | 37% | 36% | 24% |
|  | NHR | 236 | 0.36 | 0.39 | 0.34 | 0.19 | 56% | 63% | 58% | 35% |

<sup>a</sup> max2 means reducing profile to only include RFR models that correlate with assay of interest at $R^2 > 0.2$ or $R^2 > 0.05$ , whichever is better
<sup>b</sup> AA means a starting profile of all assays
<sup>c</sup> FS means a family specific starting profile
<sup>d</sup> FP means using the full profile of either the family-specific or all-assays starting profile
Note: color scale for median $R^2_{\text{ext}}$ and % $R^2_{\text{ext}} > 0.3$ are separate.

Beyond the large target classes in Table 1, there were at least some successful models from every one of the 51 smaller target sub-classes annotated in the database, indicating that pQSAR models can be used on virtually every target type. About half the models, both from NVS and ChEMBL, were phenotypic screens with no assigned target. These are particularly challenging for SA RFR models, with median *R*^2^_ext_=0.04 and only 5.0% success for NVS phenotypic assays and *R*^2^_ext_=0.09 and 24% success for ChEMBL, presumably because phenotypic assays can be modulated through a number of mechanisms. pQSAR does surprisingly well on phenotypic assays, with median *R*^2^_ext_=0.60 and 76% success for the NVS data, and median *R*^2^_ext_=0.57 and 75% success for ChEMBL.

Transfer learning between NVS and ChEMBL profiles was tested by using the ChEMBL RFR profile to build PLS models for NVS assays (Figure S4A) and *vice versa* (Figure S4B). As was previously reported for kinases, pQSAR models for NVS assays trained using the ChEMBL AA FP profile and pQSAR models for ChEMBL assays trained using the AA FP NVS profile performed little better than simple SA RFR models. Cross-dataset predictions using max2 reduced profiles are only slightly better. Combined FPs perform almost as well as native FPs. Combined max2 profiles were no better or worse than native NVS or ChEMBL max2 profiles alone. Thus, combining RFR profiles is useful for expanding the applicability domains to cover both data sets, but does not improve the accuracy over the native models.

With so many compound descriptors, chance correlations were a concern. Several Y-scrambling protocols were performed to estimate chance correlations. Y-scrambling the entire AA max2 pQSAR procedure, including building the SA RFR models on Y-scrambled IC_50_ data for each of the 11,805 assays, gave only 52 SA RFR and 47 AA max2 PLS models with *R*^2^_ext_>0.3. This is only 0.4% of the 11,805 assays. Not surprisingly, the largest of these assays contained only 81 pIC_50_s, so successful models built on larger assays are very unlikely by chance. Furthermore, 34 of those 47 “successful” AA max2 pQSAR models gave good models on real data. Given the tiny Y-scrambling rate, these are undoubtedly actual good models. One could, therefore, anticipate that only 13 out of the 8558 successful models trained on real data, *i.e.* about 0.15%, are actually chance correlations, a remarkably low false positive rate.

However, the above test is possibly too conservative, because the SA RFR matrix built on Y-scrambled data should have almost no multicollinearity. *i.e.* the rank of the SA RFR predicted IC_50_ profile matrix from Y-scrambled data is higher than that built from real data, and so has greater opportunity for chance correlations. The SA RFR profiles from the real data will still be independent of Y-scrambled training and test data in the PLS step. Three rounds of training Y-scrambled FP PLS models using the real SA RFR profiles gave only 32 to 36 models with *R*^2^_ext_>0.3. One assay was shared between each pair of repetitions, and no assays were shared among all 3, indicating that these really were chance events. Training Y-scrambled max2 PLS models using the real SA RFR profiles gave 35 models with *R*^2^_ext_>0.3. Twenty-two of the assays that gave chance correlations with scrambled data gave good correlations with real data, so this test also suggests that about 13 of the 8558 successful models, or 0.15%, are chance correlations.

Y-scrambling the entire max2 pQSAR procedure for ChEMBL gave 97 SA RFR models and 175 max2 PLS models with *R*^2^_ext_>0.3 with 112 compounds in the largest assay. 121 of those 175 “successful” models gave good models on real data, indicating 54 of the 4276 models, or 1.3% are likely due to chance. Three repetitions of the FP PLS step on Y-scrambled ChEMBL assays using the SA RFR profiles from real data gave from 84 to 92 models with *R*^2^_ext_>0.3. No assay appeared in all 3 repetitions, although from 1 to 6 were shared by pairs. Y-scrambling just the max2 PLS step of ChEMBL gave 133 models with *R*^2^_ext_>0.3. Sixty-seven of those 133 “successful” models gave good models using real data, suggesting that 66 of the 2682 are false positives. The higher 2.5% ChEMBL max2 pQSAR false positive rate, despite having half as many descriptors (*i.e.* assays), may reflect preponderance of small assays.

So far, only predictions on compounds with experimental data, that are in the SA RFR predicted IC_50_ profile table, have been presented. To make predictions on new compounds, they are first run through the SA RFR models to obtain their predicted activity profiles. The new profiles are then used in one or more PLS equations to yield the max2 pQSAR activity predictions. While the chemical space covered by the archive drifts over time, the procedure is highly automated. Every month, all assay data are downloaded from the NVS database, the data are filtered and processed, and all of the nearly 12,000 pQSAR models are rebuilt from scratch as described above. Nearly 50 billion pIC_50_s are then predicted for the 5.5 million compounds in the NVS database against the 8558 successful models. These are loaded into a Databricks database for virtual screening and general data mining. Z-scaled pIC_50_ predictions are also stored to facilitate comparison across assays, *e.g.* for predicting mechanism-of-action in phenotypic screens or potential off-target activities. The predictions are also stored in a specialized “descriptor store” database for use as continuous-valued fingerprints in clustering or other data analyses.

### Prospective application examples

Over the last 15 years pQSAR, in its several incarnations, has been applied to well over 100 projects--initially for kinases and more recently for diverse targeted and phenotypic projects. Applications have spanned drug discovery: virtual screening for tool compounds and medicinal chemistry starting points, selectivity calculation including isoform selectivity, hit list triaging, fragment hit expansion, toxicity and promiscuity prediction, mechanism-of-action prediction, virtual orthosteric counter-screens to identify allosteric inhibitors from experimental high-throughput screens, etc. A few examples below illustrate typical applications both from the original kinase pQSAR and AA FP pQSAR. Results from AA max2 projects are not yet available.

### Raf kinase virtual screening

B-Raf provides an early kinase virtual screening example. The goal was to identify potential backup scaffolds from 600K compounds added to the corporate archive since an earlier full-deck high throughput screening (HTS). Being an advanced project, over 35,000 biochemical IC_50_s and cellular EC_50_s were available divided among 5 assays: the target kinase, a mutant variant prevalent in tumors, an isoform, a cell proliferation assay and a cellular target modulation assay measuring phosphorylation of downstream kinase ERK. Compounds with predicted pIC_50_ or pEC_50_ ≥ 6 by all 5 models were initially selected. Molecular weight and rotatable bond filters were applied, and compounds containing known Raf scaffolds were removed. The remaining compounds were clustered into 47 clusters and all cluster centers, as well as analogues with Tanimoto similarity coefficient (Tc) ≥ 0.8 to the cluster centers, were kept. Seventy-three compounds were sent for biochemical evaluation. Fifty-three had IC50 ≤ 10 µM, an overall 80% hit rate. The hit-count line in the novelty/hit-rate histogram (Figure 3) for this project shows that, per usual, none of the selected compounds were from within Tc>0.8 of the training compounds. The majority of hits had similarities less than 0.6, and shared no scaffolds with the nearest known active compounds. Hit rates were 60%-80% even for compounds with novel scaffolds and similarity Tc < 0.4 to any of the 35K known actives, 2 logs above the HTS background rate.

**Figure 3.**
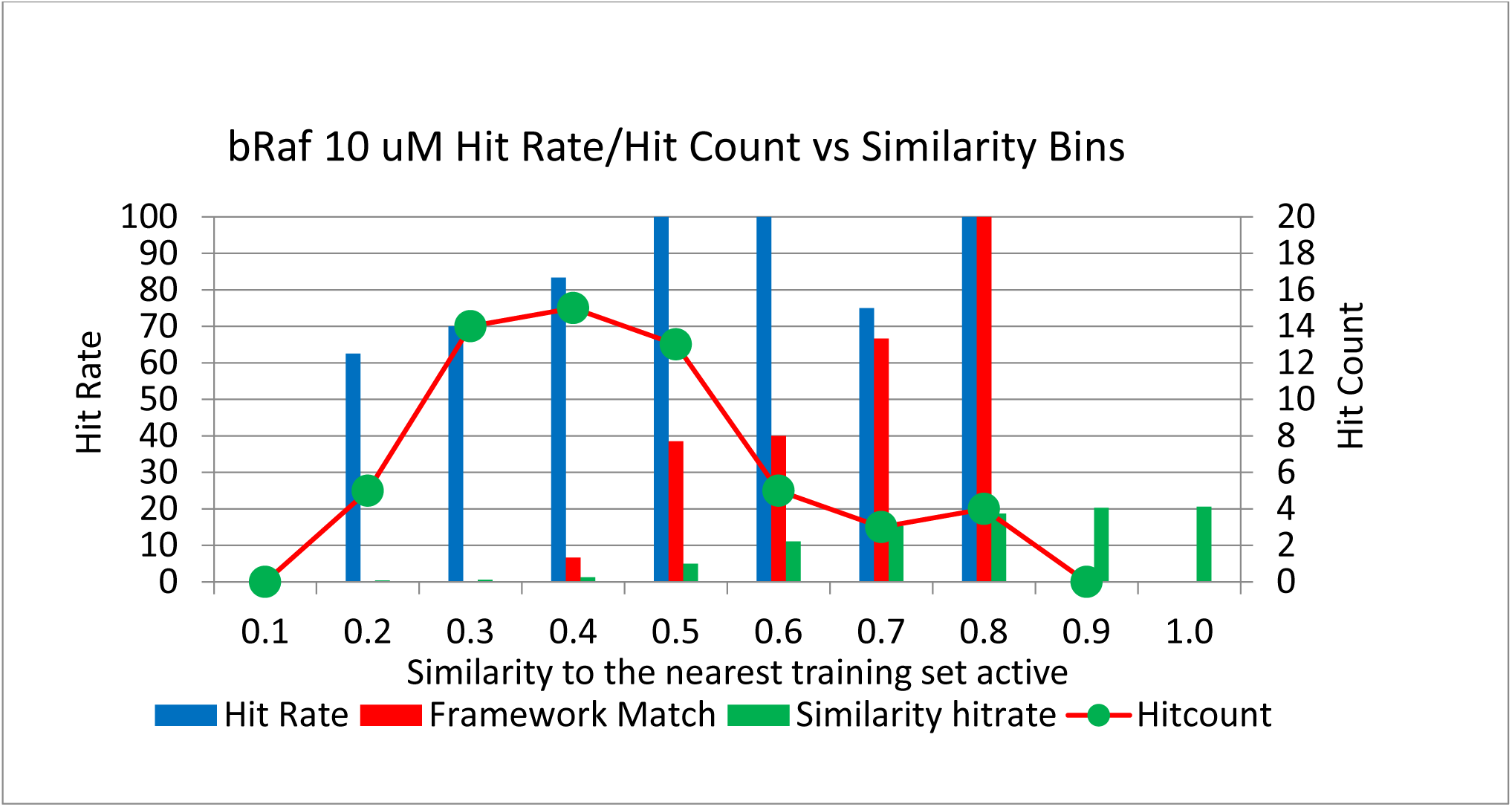
A novelty/hit-rate histogram for bRaf with similarity bins on the X-axis, hit rate on the primary Y-axis and hit count on the secondary Y-axis. The blue bars represent hit rates from the virtual screen, while the red bars plot the percentage of hits in each bin that share a Bemis & Murcko framework with a framework in the training set. The green circles show the hit counts for each similarity bin. Green bars show hit rate from an earlier 10 µM HTS binned by similarity. No compounds were selected from the bins most similar to the training set.

### Understanding RAF cellular assays

The Raf project also supplies an example for cellular assays. While interpreting them can potentially be misleading due to multi-collinearities, pQSAR PLS coefficients can carry useful information about mechanisms-of-action. The target modulation (TM) and cell proliferation (CP) assays differed for A375 cells with bRAFV599E, a mutant isoform over expressed in melanoma cell lines. While biochemical bRAFV999E IC_50_s trended with the TM EC_50_s, with *R*^2^ = 0.44 for 2302 compounds, the correlation with the CP EC_50_s in the same A375 cell line was lower, with *R*^2^ = 0.24 for 1657 compounds. The target modulation assay measured inhibition of a phosphorylation endpoint in the RAF/MEK/ERK pathway, believed to be specific to that signaling cascade. Although the cell line was described as “driven” by mutant bRAF, cell proliferation can be modulated by multiple mechanisms. To look for other interfering kinases, pQSAR models were built for both the A375 TM and A375 CP assays using only a profile of 92 kinase biochemical models, without including cellular assays. The PLS coefficients for the TM and CP pQSAR models are plotted on the Sugen kinome dendrograms in Figure S5. The diameters of the circles indicate the magnitudes of the PLS coefficients, blue indicating a positive contribution and red negative. For the bRAF TM model, the 2 largest positive coefficients were bRAF (relative value = 1) and cRAF (relative value = 0.54), consistent with target modulation being driven primarily by RAF inhibition. In the cell proliferation assay, bRAF is still most important (relative value = 0.86), but some other tyrosine kinases (TK) also had large positive coefficients. PDGFRb (relative value = 0.71) was selected for further analysis. For many of the compounds inactive at the highest tested concentration of 10 µM (pIC_50_ < 5) in the CP assay, bRAFV599E biochemical pIC_50_ was high, but in these cases PDGFRb pIC_50_ < 6.5. Conversely, when A375 CP pEC_50_ > 5.5, both bRAFV599E and PDGFRb were above pIC_50_ ~ 7. Thus, the CP assay was most sensitive to bRAF/PDGFRb dual inhibitors. PDGFRs were reported to activate Ras via SHP2, making this interactions plausible ^21^.

This analysis was done in 2009. In 2010, a publication by Murphy *et al.* ^22^ reporting a selective dual PDGFRb/RAF inhibitor efficacious in murine models of pancreatic and kidney tumors supported the interplay between these two kinases. A publication in December of 2010 by Nazarian *et al.*^*23*^ and a follow-up publication by Solit *et al.*^*24*^ in January of 2011 described PLX-4032, a RAF inhibitor then in clinical trials for melanoma, showing resistance and tumor regrowth in patients six months into treatment. Analysis showed that PDGFRb was up-regulated in a number of the clinical tumor samples. They hypothesized that the tumors developed resistance to RAF inhibitor treatment and shutdown of the MAPK pathway by up-regulating the PDGFRb driven pathway. The follow-up publication suggested co-treatment with RAF and PDGFRb inhibitors as a possible way of overcoming this resistance. Analyzing the pQSAR coefficients and data had anticipated these results later reported from the clinic. This suggests that pQSAR coefficients capture underlying biochemical mechanisms, not mere meaningless curve fitting, and can be used to identify mechanisms-of-action.

### Kinase host off-target prediction from malaria phenotypic screening hits

Attempting to predict potential human anti-targets for a series of malaria inhibitors, a pQSAR model was trained on malaria phenotypic EC_50_s using a profile of 192 human kinase biochemical assays. 3 months later, inhibition of human haspin (GSG2), the kinase with the second largest PLS coefficient, was experimentally confirmed in profiling by DiscoverX, again indicating that the coefficients of pQSAR models capture real biochemical mechanisms.

### SGK3 kinase virtual selectivity screen

An SGK3 example illustrates the use of kinase pQSAR in predicting off-target activity. A tool compound to help validate SGK3 as a drug target required broad kinase selectivity, as AGC, TK and PI3K kinases were expected to interfere with the target modulation cellular assay. 100,000 compounds from a “kinase focused” deck were screened experimentally at a single 10 µM concentration. About 4,000 inhibited SGK3 at greater than 30%. Median predicted TK IC_50_ greater than 1 µM, and AGC IC_50_ greater than 10 µM from 28 TK kinase and 10 AGC kinase pQSAR models were used as selectivity filters. Cellular exposure was estimated from the median ratio of predicted cellular EC_50_ and biochemical IC_50_ (B/TM) < 0.3 for 13 pairs of kinase target modulation pQSAR models. Using these filters and standard drug-like requirements, 750 compounds were selected for confirmation in an 8-concentration SGK3 IC_50_ assay. Sixty compounds had IC_50_ < 1 µM while 330 had IC_50_ < 10 µM.

An SGK3 pQSAR model built from the 750 IC_50_s, with a modest *R*^2^_ext_ = 0.32, was used to select and screen an additional 567 compounds from the archive. A total of 327 had IC_50_ < 10 µM, a 58% hit rate. The novelty/hit-rate histogram in Figure S6 shows hit rates of 40% - 60% even for compounds with Tc < 0.5 to the training set models. With so many novel medchem starting points at hand, a conventional HTS was canceled, and the team instead began developing a more relevant but complex cascade assay.

A second round of off-target filtering selected 30 compounds with experimental SGK3 IC_50_ < 5 µM, and that were selective based on the pQSAR predictions and available experimental IC_50_s, including at least 2 experiments from AGC, TK or PI3K kinases. Twenty of these 30 compounds, hand selected by experienced medicinal chemists, were sent for biochemical profiling against a panel of 70 kinases. The best of these had SGK3 IC_50_ = 150 nM, and IC_50_ > 1 µM against the panel. This compound had excellent physical properties: CaCO-2 A-B=3, B-A=2, solubility = 10 M, and ClogD at pH 7.4 = 0.44. Thus, the kinase-family pQSAR models had rapidly identified a selective, cell permeable compound from the experimental MTS that met the SGK3 team’s criteria for a tool compound.

### An epigenetic target

The NVS archive was virtually screened with AA FP pQSAR models trained on 3 assays of slightly different formats for an epigenetic target. About 9000 compounds predicted to be active in any of the 3 models were selected. Compounds with Tc>0.8 to any known sub-µM compounds or with undesired functionality were removed. The remaining ~5500 compounds were binned for predicted high potency, high ligand efficiency or high lipid efficiency. AA FP pQSAR predicted selectivity over 2 other epigenetic targets were combined with predicted on-target activity into one score. Compounds from each bin were clustered and representative compounds of each cluster with good scores were selected. The final collection of 1003 unique compounds was experimentally tested in a single-dose, percentage inhibition, biochemical assay. A total of 109 compounds showing at least 30% inhibition were considered as hits and sent to the next validation step. The novelty/hit-rate histogram in Figure S7 shows that about 70% of the hits were highly novel, coming from bins of Tc=0.2 and 0.3 to the known inhibitors.

### Malaria phenotypic virtual screen

An AA FP pQSAR model was trained on 9,822 IC_50_s from an experimental phenotypic malaria screen. 22,546 predicted actives were selected based on novelty, potency, ligand efficiency and drug-like properties. An additional complement diversity selection of 34,419 was designed to sample the remainder of the archive. 94% of the 5,636 hits with confirmed IC_50_<10uM, and 98% of the 880 hits with IC_50_<100nM came from the pQSAR arm. Figure S8 shows that 75% of the confirmed pQSAR hits had novel scaffolds and nearly 2,000 had Tc<0.6 to the known inhibitors.

### Practical considerations

While pQSAR is not the only multi-task method, it does have some practical strong points. It is simple, embarrassingly parallel and scales well over a conventional cluster. There are no hyper-parameters to retrain even when scaling from a dozen to 10,000 assays. Unlike conventional multi-task deep neural networks (MT-DNN), it allows for a separate training/test set split for each assay, important for realistic model evaluation. Unlike MT-DNNs or matrix imputation methods, including the recently published Alchemite method^25^, it allows a separate subset of tasks for each assay, *e.g.* max2, reducing unwanted noise. Our current implementation uses simple SA RFR on Morgan fingerprints, followed by PLS regression. However, one can easily replace or augment these 2D RFR models by additional 2D or 3D QSAR models, pharmacophore models or docking models in the first step, for a richer description of the chemistry. Similarly, additional column types such as physicochemical properties or genomics data can be added beside the assays for a richer description in the experimental dimension. Thus, there are myriad opportunities to extend the scope of pQSAR beyond this basic beginning.

## CONCLUSIONS

pQSAR for single protein families was previously shown to far exceed single-assay QSAR models. All assay pQSAR is a unified platform that included all of the 51 annotated target sub-classes, as well as 5506 phenotypic assays, which expands the applicability domain for idiosyncratic targets or phenotypic screens. AA max2 profile reduction further removes noise and enhances model performance. AA max2 also saves computation. The number of PLS descriptors was reduced by about 80% and 96% at the thresholds of *R*^2^=0.05 and *R*^2^=0.20, respectively, while achieving improved performance. This is a substantial computational resource saving when training 11,805 PLS models and predicting 5.5 million compounds on each. The negligible chance correlations indicate the reliability of AA max2 pQSAR models.

Nearly 50 billion pIC_50_s are predicted for the 5.5 million compounds in the NVS database against the 8558 successful models. The pQSAR models are updated monthly. As newly tested dose-response pIC_50_s are continuously adding to NVS database, the number of successful models as well as the predicted pIC_50_s are expected to grow rapidly.

## SUPPORTING INFORMATION

The curated pIC_50_s of 0.5 million compounds measured on 4276 ChEMBL assays as well as their prediction can be found in the supplementary materials.

**Table S1**. General information of 4276 ChEMBL assays. The specific assay identifiers, the number of tested compounds, corresponding assay type and target family. (TXT)

**Table S2**. The ChEMBL ID, associated assay, observed pIC_50_s and predicted pIC_50_s by max2 PLS models for 1.4 million activity values. (TXT)

**Table S3**. The ChEMBL ID and SMILES of half million unique ChEMBL compounds. (CSV)

**Figure S1**. Histogram of the number of tested compounds in each of 11,805 NVS assays (A) and 4,276 ChEMBL assays (B). (PDF)

**Figure S2.** Box plots summarizing historgrams of Tanimoto similarity between test set compounds and their nearest neighbors in the training sets. A: “realistic” split and B: random split for NVS data. C: “realistic” split and D: random split for ChEMBL data. (PDF)

**Figure S3**. Comparison of variable selection between recursive random forest regression (RRFR) and PLS-based pQSAR. A: The median *R*^2^_ext_ for NVS RRFR models is 0.51 and 7317 of them with *R*^2^_ext_ > 0.3. This is slightly better than (PLS) FP pQSAR, but worse than (PLS) max2 pQSAR. B: the median *R*^2^_ext_ for ChEMBL RRFR models is 0.29 and 2114 of them with *R*^2^_ext_ > 0.3. (PDF)

**Figure S4**. Cross prediction performance for pQSAR models for all NVS assays trained on the ChEMBL RFR profile (A) and all ChEMBL assays trained on the NVS profile (B) perform little better than SA RFR models. Both NVS and ChEMBL assays are only slightly better predicted by each other’s max2 profiles. Combined FPs perform almost as well as native FPs. Combined max2 profiles showed no advantage or disadvantage over native ChEMBL max2 or NVS max2 profiles alone. All evaluations are for the realistic test sets. FP: full profile; NVS&ChEMBL: combined NVS and ChEMBL profiles; max2: The maximum *R*^2^_ext_ using thresholds of *R*^2^>0.05 and *R*^2^>0.2. (PDF)

**Figure S5**. Sugen kinome trees showing kinase pQSAR PLS coefficients of biochemical kinase assays for the (a) target modulation and (b) cell proliferation cellular RAF assays. The size represents the magnitude of the coefficient, blue being positive and red being negative. (PDF)

**Figure S6**. A novelty/hit-rate histogram for SGK3 with similarity bins on the X-axis, hit rate on the primary Y-axis and hit count on the secondary Y-axis. The blue bars represent hit rates from the virtual screen, while the orange bars plot the percentage of hits in each bin which share a Bemis & Murcko framework with a framework in the training set. The grey circles show the hit counts for each similarity bin.. (PDF)

**Figure S7**. A novelty/hit-rate histogram for an epigenetic target with similarity bins on the X-axis, hit rate on the primary Y-axis and hit count on the secondary Y-axis. The blue bars represent hit rates from the virtual screen, while the red bars plot the percentage of hits in each bin which share a Bemis & Murcko framework with a framework in the training set. The green circles show the hit counts for each similarity bin.

**Figure S8**. A novelty/hit-rate histogram for malaria phenotypic virtual screen. Red bars are total confirmed pQSAR hits, and blue bars are the number of those with novel scaffolds. Yellow and green bars are total and novel hits from a compliment diversity selection. (PDF)

## Notes

The authors declare no competing financial interest.

### ACKNOWLEDGEMENTS

Xin Liu would like to thank the Novartis Young Entrepreneur Program 2017 (NYEP) for funding. Zhengtian Yu and Miao Dai are very much appreciated for their help and efforts for Xin Liu in this program and in the extended internship. The authors express their gratitude to the thousands of medicinal chemists and assay biologists that worked on all NVS assays over many decades that represent the real effort behind the success of pQSAR models.

## ABBREVIATIONS

AA: all assay
CP: cell proliferation
FP: full profile
FS: family-specific
GPCR: G-protein coupled receptor
HTS: high throughput screening
max2: reducing profile to only include RFR models that correlate with assay of interest at *R*^2^>0.2 or *R*^2^>0.05, whichever is better.
MT-DNN: multi-task deep neural networks
NHR: nuclear hormone receptor
NVS: Novartis
pIC_50_s: the negative logarithm of IC_50_s
PLS: partial least squares
pQSAR: profile-QSAR
QSAR: quantitative structure-activity relationship
*R*^2^_ext_: the coefficient of determination of external realistic test set.
RFR: random forest regression
RP: reduced profile
RRFR: recursive random forest regression
SA: single assay
Tc: Tanimoto similarity coefficient
TK: tyrosine kinases
TM: target modulation

## For Table of Contents Use Only

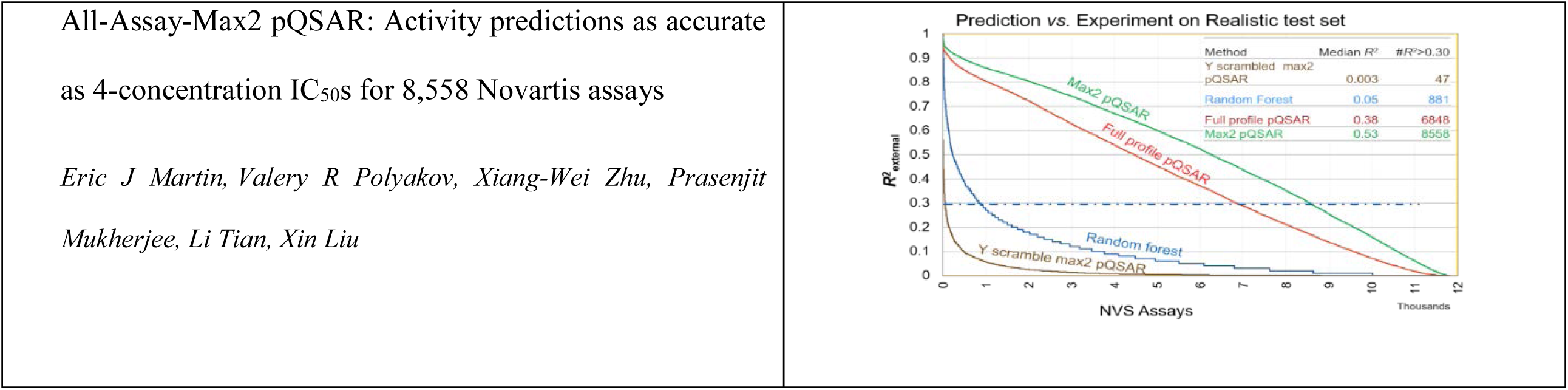

## Notes

#### Summary of Updates

Added training/test set split indication to file S2

## REFERENCES

1. Tropsha, A.; Golbraikh, A., Predictive QSAR Modeling Workflow, Model Applicability Domains, and Virtual Screening. Current Pharmaceutical Design 2007, 13, 3494–3504.

2. Ramsundar, B. K., Steven M.; Riley, Patrick; Webster, Dale; Konerding, David E: Pande, Vijay S, Massively Multitask Networks for Drug Discovery. ArXiv 2015, 1502.02072.

3. Kauvar, L. M.; Higgins, D. L.; Villar, H. O.; Sportsman, J. R.; Engqvist-Goldstein, Å.; Bukar, R.; Bauer, K. E.; Dilley, H.; Rocke, D. M., Predicting ligand binding to proteins by affinity fingerprinting. Chemistry & Biology 1995, 2, 107–118.

4. Zhu, H.; Rusyn, I.; Richard, A.; Tropsha, A., Use of cell viability assay data improves the prediction accuracy of conventional quantitative structure-activity relationship models of animal carcinogenicity. Environ Health Persp 2008, 116, 506–513.

5. Russo, D. P.; Kim, M. T.; Wang, W. Y.; Pinolini, D.; Shende, S.; Strickland, J.; Hartung, T.; Zhu, H., CIIPro: a new read-across portal to fill data gaps using public large-scale chemical and biological data. Bioinformatics 2017, 33, 464–466.

6. Barycki, M.; Sosnowska, A.; Jagiello, K.; Puzyn, T., Multi-Objective Genetic Algorithm (MOGA) As a Feature Selecting Strategy in the Development of Ionic Liquids’ Quantitative Toxicity– Toxicity Relationship Models. Journal of Chemical Information and Modeling 2018, 58, 2467–2476.

7. Russo, D. P.; Strickland, J.; Karmaus, A. L.; Wang, W.; Shende, S.; Hartung, T.; Aleksunes, L. M.; Zhu, H., Nonanimal Models for Acute Toxicity Evaluations: Applying Data-Driven Profiling and Read-Across. Environ Health Persp 2019, 127, 47001–47001.

8. Martin, E.; Mukherjee, P.; Sullivan, D.; Jansen, J., Profile-QSAR: a novel meta-QSAR method that combines activities across the kinase family to accurately predict affinity, selectivity, and cellular activity. J Chem Inf Model 2011, 51, 1942–56.

9. Martin, E. J.; Polyakov, V. R.; Tian, L.; Perez, R. C., Profile-QSAR 2.0: Kinase Virtual Screening Accuracy Comparable to Four-Concentration IC_50_s for Realistically Novel Compounds. Journal of Chemical Information and Modeling 2017, 57, 2077–2088.

10. Mukherjee, P.; Martin, E., Profile-QSAR and Surrogate AutoShim Protein-Family Modeling of Proteases. J. Chem. Inf. Model. 2012, 52, 2430–2440.

11. Tian, L. M., Eric, Exploring protein families with Profile-QSAR. American Chemical Society: Washington, DC, 2015.

12. Baumann, K.; Stiefl, N., Validation tools for variable subset regression. J Comput Aid Mol Des 2004, 18, 549–562.

13. Gaulton, A.; Hersey, A.; Nowotka, M.; Bento, A. P.; Chambers, J.; Mendez, D.; Mutowo, P.; Atkinson, F.; Bellis, L. J.; Cibrián-Uhalte, E.; Davies, M.; Dedman, N.; Karlsson, A.; Magariños, M. P.; Overington, J. P.; Papadatos, G.; Smit, I.; Leach, A. R., The ChEMBL database in 2017. Nucleic acids research 2017, 45, D945–D954.

14. Butina, D., Unsupervised Data Base Clustering Based on Daylight’s Fingerprint and Tanimoto Similarity: A Fast and Automated Way To Cluster Small and Large Data Sets. Journal of Chemical Information & Modeling 1999, 39, 747–750.

15. Pedregosa, F.; Varoquaux, G.; Gramfort, A.; Michel, V.; Thirion, B.; Grisel, O.; Blondel, M.; Prettenhofer, P.; Weiss, R.; Dubourg, V.; Vanderplas, J.; Passos, A.; Cournapeau, D.; Brucher, M.; Perrot, M.; Duchesnay, E., Scikit-learn: Machine Learning in {P}ython. J Mach Learn Res 2011, 12, 2825–2830.

16. Rücker, C.; Rücker, G.; Meringer, M., y-Randomization and Its Variants in QSPR/QSAR. Journal of Chemical Information and Modeling 2007, 47, 2345–2357.

17. Clark, M.; Cramer Iii, R. D., The Probability of Chance Correlation Using Partial Least Squares (PLS). Quantitative Structure-Activity Relationships 1993, 12, 137–145.

18. Zhu, X.-W.; Xin, Y.-J.; Ge, H.-L., Recursive random forests enable better predictive performance and model interpretation than variable selection by LASSO. Journal of Chemical Information and Modeling 2015, 55, 736–746.

19. Lavecchia, A., Machine-learning approaches in drug discovery: methods and applications. Drug Discov Today 2015, 20, 318–331.

20. Fernandez-Delgado, M.; Cernadas, E.; Barro, S.; Amorim, D., Do we Need Hundreds of Classifiers to Solve Real World Classification Problems? J Mach Learn Res 2014, 15, 3133–3181.

21. Agazie, Y. M.; Hayman, M. J., Molecular mechanism for a role of SHP2 in epidermal growth factor receptor signaling.

22. Murphy, E. A.; Shields, D. J.; Stoletov, K.; Dneprovskaia, E.; McElroy, M.; Greenberg, J. I.; Lindquist, J.; Acevedo, L. M.; Anand, S.; Majeti, B. K.; Tsigelny, I.; Saldanha, A.; Walsh, B.; Hoffman, R. M.; Bouvet, M.; Klemke, R. L.; Vogt, P. K.; Arnold, L.; Wrasidlo, W.; Cheresh, D. A., Disruption of angiogenesis and tumor growth with an orally active drug that stabilizes the inactive state of PDGFRbeta/B-RAF. Proceedings of the National Academy of Sciences of the United States of America 2010, 107, 4299–4304.

23. Nazarian, R.; Shi, H. B.; Wang, Q.; Kong, X. J.; Koya, R. C.; Lee, H.; Chen, Z. G.; Lee, M. K.; Attar, N.; Sazegar, H.; Chodon, T.; Nelson, S. F.; McArthur, G.; Sosman, J. A.; Ribas, A.; Lo, R. S., Melanomas acquire resistance toB-RAF(V600E) inhibition by RTK or N-RAS upregulation. Nature 2010, 468, 973–U377.

24. Solit, D. B.; Rosen, N., Resistance to BRAF inhibition in melanomas. N Engl J Med 2011, 364, 772–4.

25. Whitehead, T. M.; Irwin, B. W. J.; Hunt, P.; Segall, M. D.; Conduit, G. J., Imputation of Assay Bioactivity Data Using Deep Learning. Journal of Chemical Information and Modeling 2019.

