## Supplemental Figures for "All-Assay-Max2 pQSAR: Activity predictions as accurate as 4-concentration IC_50_s for 8,558 Novartis assays"

Figure S1

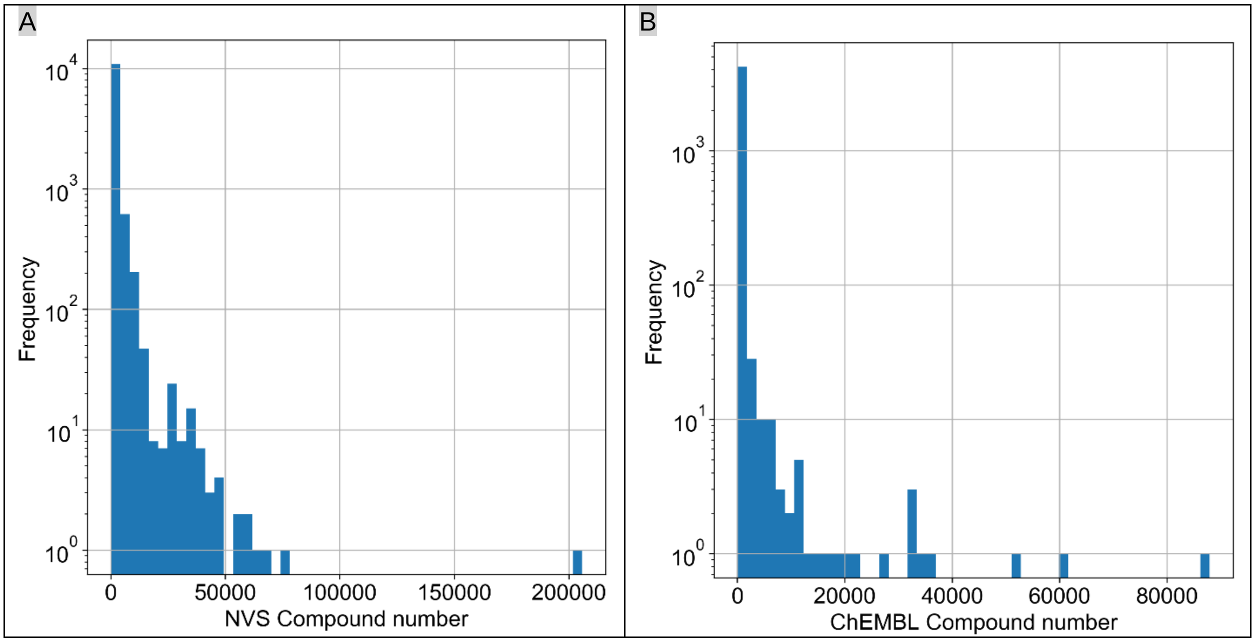


**Figure S1**. Histogram of the number of tested compounds in each of 11,805 NVS assays (A) and 4,276 ChEMBL assays (B).

Figure S2

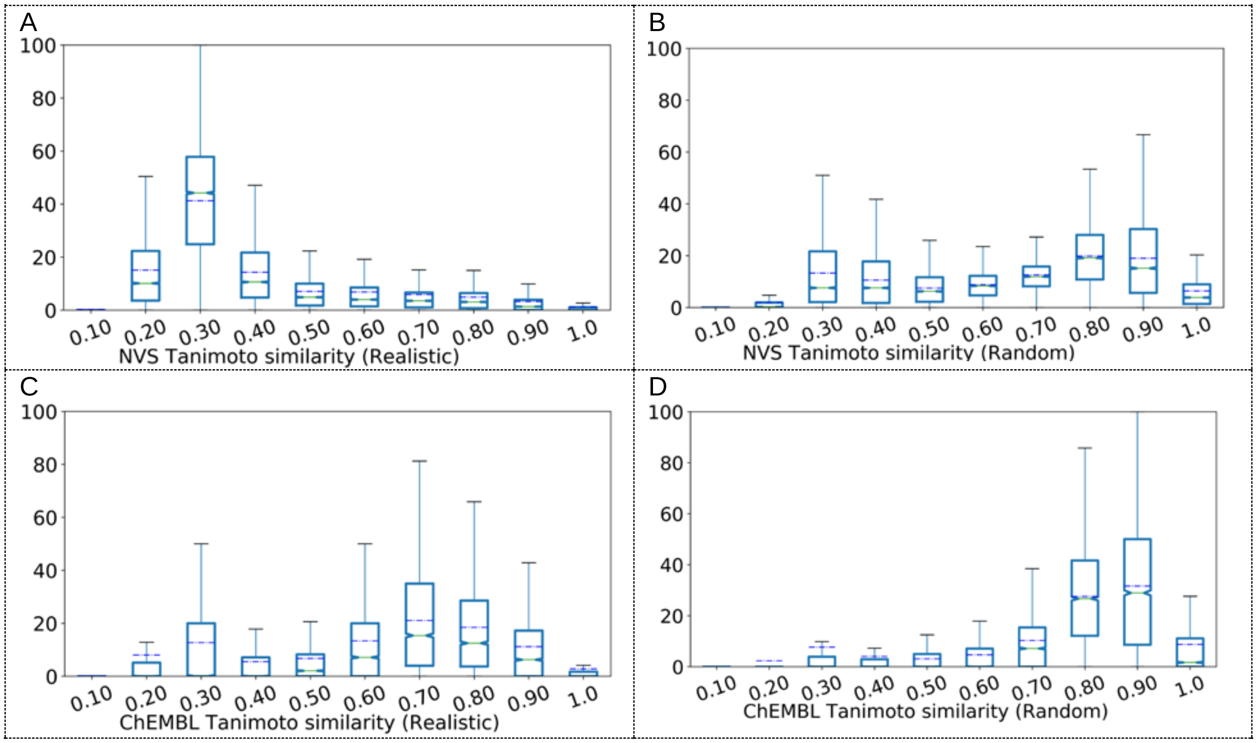


**Figure S2**. Box plots summarizing historgrams of Tanimoto similarity between test set compounds and their nearest neighbors in the training sets. A: “realistic” split and B: random split for NVS data. C: “realistic” split and D: random split for ChEMBL data.

Figure S3.

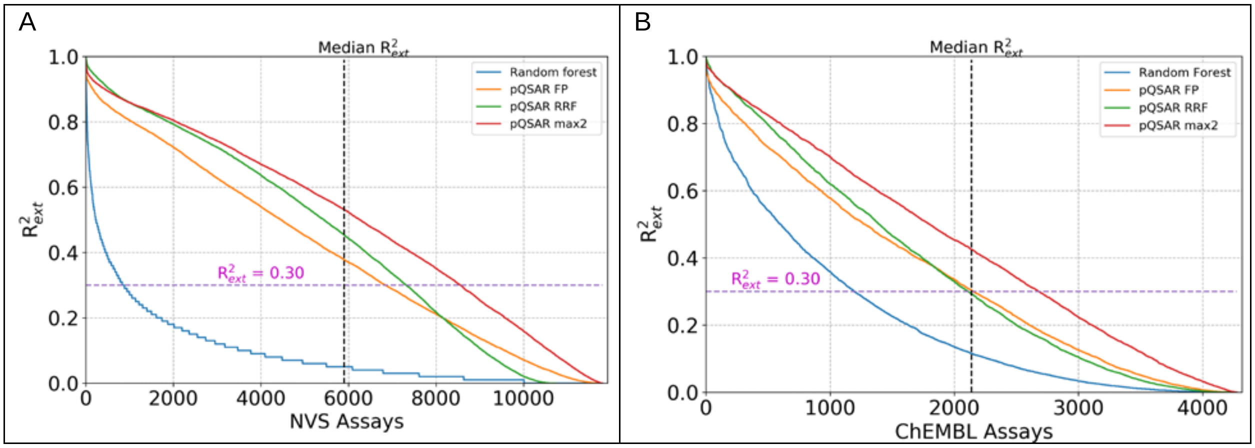


**Figure S3**. Comparison of variable selection between recursive random forest regression (RRFR) and PLS-based pQSAR. A: The median $R^2_{\mathrm{ext}}$ for NVS RRFR models is 0.51 and 7317 of them with $R^2_{\mathrm{ext}} > 0.3$. This is slightly better than (PLS) FP pQSAR, but worse than (PLS) max2 pQSAR..B: the median $R^2_{\mathrm{ext}}$ for ChEMBL RRFR models is 0.29 and 2114 of them with $R^2_{\mathrm{ext}} > 0.3$.

Figure S4

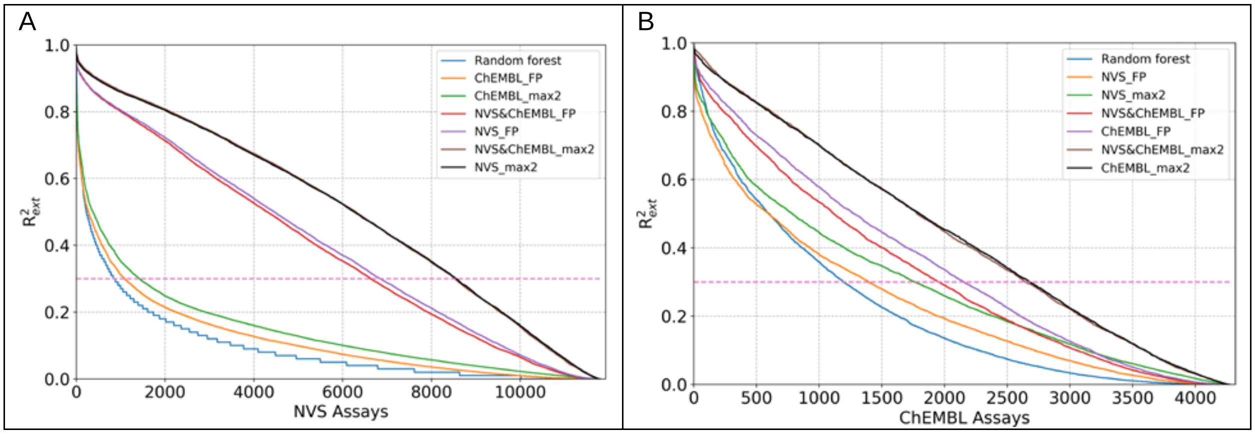


**Figure S4**. Cross prediction performance for pQSAR models for all NVS assays trained on the ChEMBL RFR profile (A) and all ChEMBL assays trained on the NVS profile (B) perform little better than SA RFR models. Both NVS and ChEMBL assays are only slightly better predicted by each other's max2 profiles. Combined FPs perform almost as well as native FPs. Combined max2 profiles showed no advantage or disadvantage over native ChEMBL max2 or NVS max2 profiles alone. All evaluations are for the realistic test sets. FP: full profile; NVS&ChEMBL: combined NVS and ChEMBL profiles; max2: The maximum $R^2_{ext}$ using thresholds of $R^2$>0.05 and $R^2$>0.2.

Figure S5

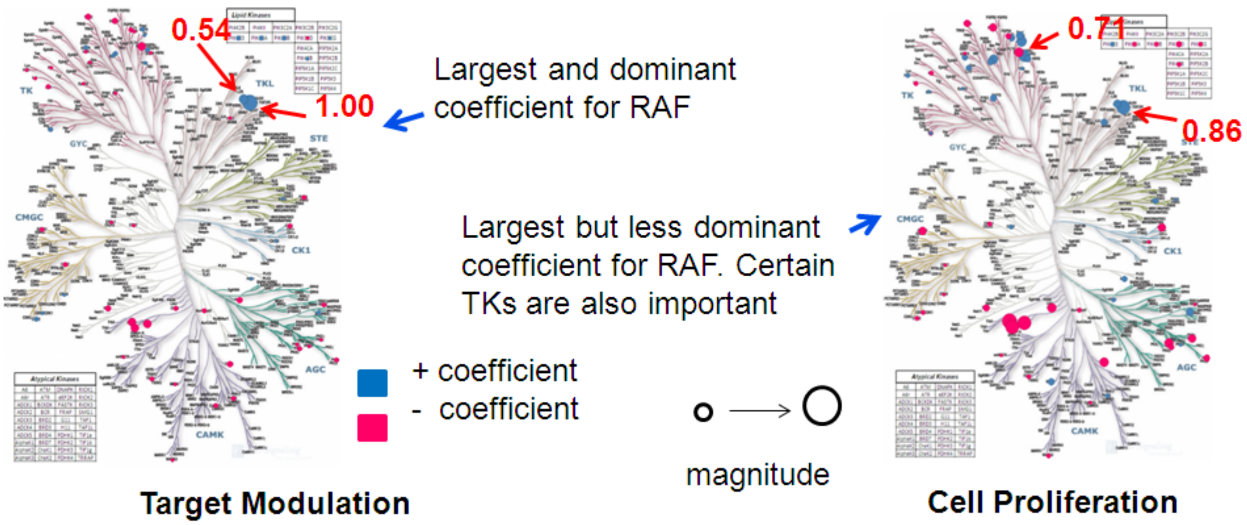


**Figure S5**. Sugen kinome trees showing kinase pQSAR PLS coefficients of biochemical kinase assays for the (a) target modulation and (b) cell proliferation cellular RAF assays. The size represents the magnitude of the coefficient, blue being positive and red being negative.

Figure S6

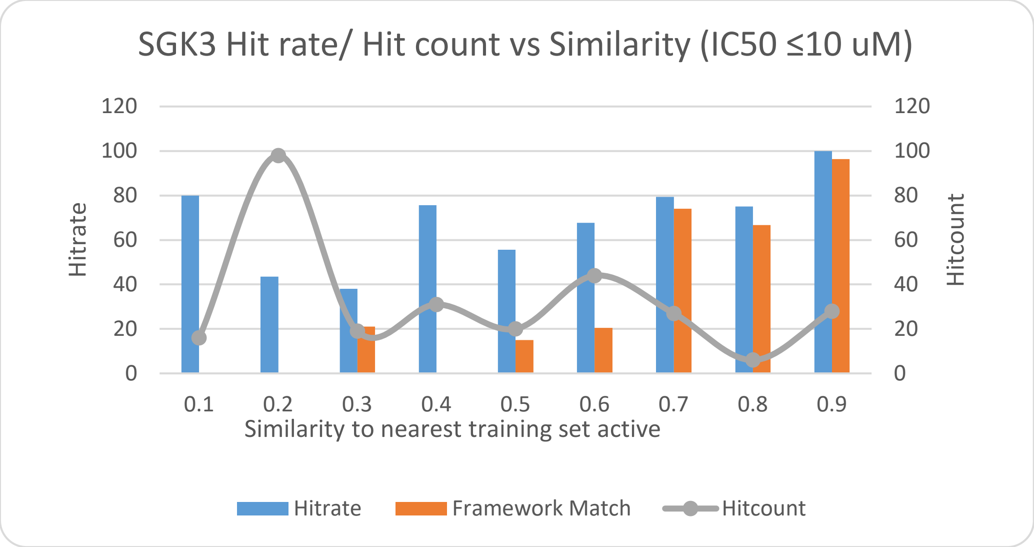


**Figure S6.** A novelty/hit-rate histogram for SGK3 with similarity bins on the X-axis, hit rate on the primary Y-axis and hit count on the secondary Y-axis. The blue bars represent hit rates from the virtual screen, while the orange bars plot the percentage of hits in each bin which share a Bemis & Murcko framework with a framework in the training set. The grey circles show the hit counts for each similarity bin.

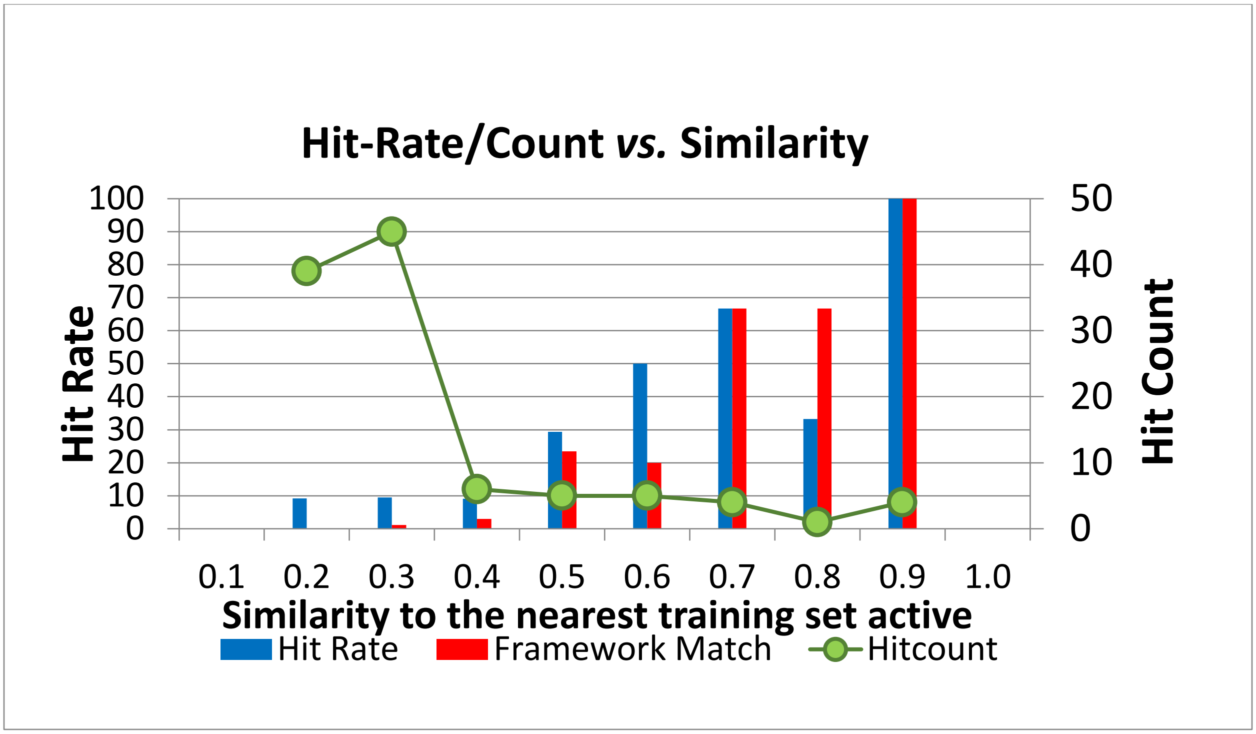


**Figure S7**. A novelty/hit-rate histogram for an epigenetic target with similarity bins on the X-axis, hit rate on the primary Y-axis and hit count on the secondary Y-axis. The blue bars represent hit rates from the virtual screen, while the red bars plot the percentage of hits in each bin which share a Bemis & Murcko framework with a framework in the training set. The green circles show the hit counts for each similarity bin.

Figure S7

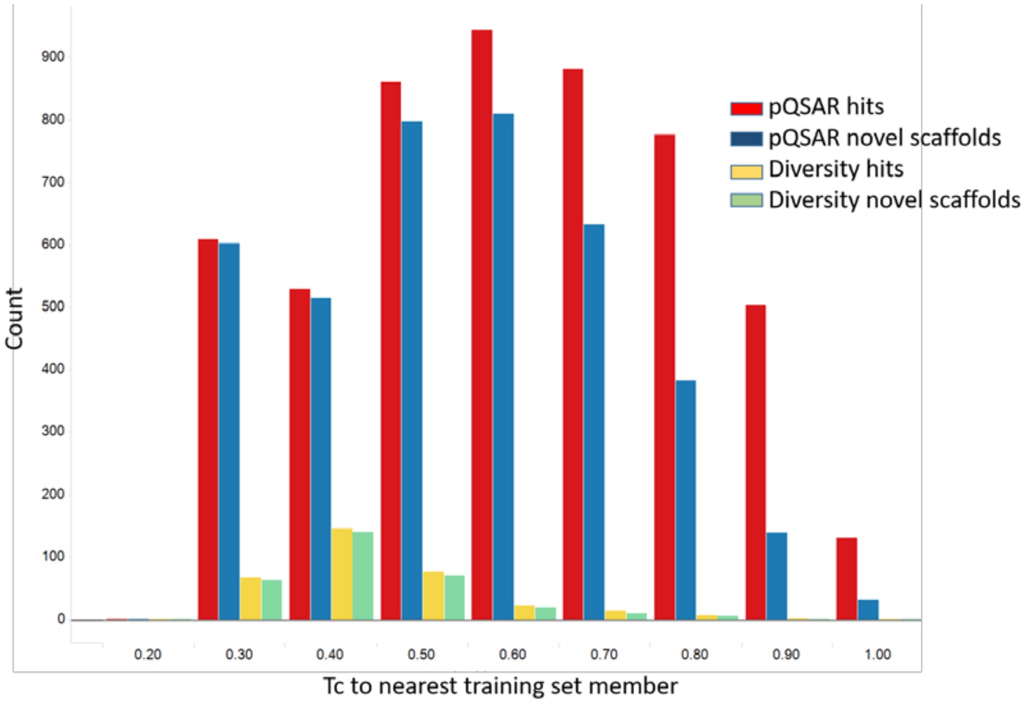


**Figure S8**. A novelty/hit-rate histogram for the malaria phenotypic virtual screen. Red bars are total confirmed pQSAR hits, and blue bars are the number of those with novel scaffolds. Yellow and green bars are total and novel hits from a compliment diversity selection.
